# ProChoreo: *de novo* Binder Design from Conformational Ensembles with Generative Deep Learning

**DOI:** 10.64898/2026.01.23.701298

**Authors:** Saisai Ding, Yi Zhang

## Abstract

Deep learning has transformed protein structure prediction and *de novo* protein design; however, most existing frameworks operate on a single static conformation and underutilize the conformational heterogeneity that governs protein binding and function. We introduce ProChoreo, a generalizable framework for *de novo* binder design that explicitly incorporates conformational ensembles. ProChoreo is pretrained with multimodal contrastive learning to align protein sequences with corresponding molecular dynamics (MD)-derived ensembles, producing a shared latent representation that captures both sequence-level and dynamic structural information. This representation is then integrated into an autoregressive generator to design protein binders conditioned on receptor sequences. Designed binders are evaluated using Boltz 1 for complex structure and interaction quality, followed by MD simulations of complexes with two representative receptors: the human sweet taste receptor TAS1R2 and FGFR2. ProChoreo designs binders that encode conformational features, highlighting dynamics-informed design as a route to protein design.

## Introduction

Recent advances in deep learning have profoundly transformed the field of structural biology, providing powerful frameworks for predicting and designing protein structures. Models such as AlphaFold^1^, and RoseTTAFold^2^ employ various neural network architectures to infer three-dimensional conformations with accuracy comparable to experimental methods, markedly reducing the time from amino acid sequence to structural analysis. Structure-based deep learning frameworks such as RFdiffusion^3^, ProteinMPNN^4^, and BindCraft^5^ have demonstrated remarkable capabilities in *de novo* protein design. These models design sequences compatible with a single static backbone conformation, enabling the rational engineering of proteins with precisely defined tertiary structures. In therapeutic contexts, such approaches have been successfully applied to design high-affinity binders targeting clinically relevant receptors, including Structure-based deep learning frameworks such as Programmed cell death protein 1 (PD-1) and epidermal growth factor receptor (EGFR), achieving nanomolar binding affinities and demonstrating the feasibility of structure-conditioned binder generation^6^. Beyond binding, RFdiffusion has been used to create *de novo* hydrolases exhibiting catalytic efficiencies up to 2.2 × 10^5^ M−1 s^−1^, with folds distinct from natural serine hydrolases^7^. In parallel, sequence-based generative models, such as ESM2^8^ and ESM3^9^, extend the principles of language modeling to protein design, learning contextual relationships within amino acid sequences without explicit structural input. Notably, ESM3 has been used to generate and experimentally validate fluorescent proteins, identifying a variant with bright fluorescence that shares only 58 % sequence identity with any known fluorescent protein^9^.

Proteins are not rigid entities; instead, they populate ensembles of interconverting conformational states that are fundamental to recognition, catalysis, and regulation^10-12^. Using AF-Cluster, researchers identified and experimentally validated multiple states of KaiB and Mpt53, underscoring the biological significance of conformational diversity^13^. To computationally capture such dynamics, recent advances employ machine learning force fields (MLFFs) such as AI^2^BMD^14^ and DeepPotential^15^, which achieve accuracy similar to density functional theory in force estimation at greatly reduced cost, as well as diffusion-based and flow-based frameworks including AlphaFLOW^16^, ESMFLOW^16^, and MDGEN^17^, which extend architectures from video generation to efficiently produce molecular dynamics trajectories and plausible conformational ensembles^18,19^. In parallel, BioEmu generates protein equilibrium ensembles with generative deep learning^20^. Yet despite these developments, structure-based design frameworks remain constrained to static conformations, and generative approaches that account for conformational ensembles have not been incorporated into protein design, which is a critical limitation motivating the present study.

A promising direction to bridge this gap is the application of multimodal contrastive learning, inspired by frameworks such as CLIP^21^, which align heterogeneous modalities in a joint embedding space. In our context, this paradigm is first employed as a pretraining objective to align protein sequences with their corresponding conformational ensembles derived from molecular dynamics simulations. By jointly embedding these two modalities, the model learns a shared latent representation that captures both sequence level and dynamic structural information. This aligned space then serves as the foundation for a downstream autoregressive generator, which designs protein binders conditioned on receptor sequences.

In this study, we introduce ProChoreo, a generalizable framework for *de novo* binder generation informed by conformational dynamics. We first pretrain the model through contrastive learning between protein sequences and conformational ensembles, aligning the latent representations of both modalities to capture the intrinsic relationship between amino acid composition and conformational variability. The resulting fused embedding, distilled from this pretraining stage, is then integrated into an autoregressive generator to design proteins that explicitly encode structural features derived from conformational ensembles. To evaluate the designed binders, we employ the open-source implementations of Boltz-1 for structural and interaction assessment^22^. Finally, molecular dynamics simulations are performed on complexes formed between the designed binders and two representative receptors, human sweet taste receptor type 1 member 2 (TAS1R2) and fibroblast growth factor receptor 2 (FGFR2), to estimate binding affinities and validate the functional relevance of the generated sequences. Together, these results illustrate the potential of integrating conformational ensemble information into binder design frameworks.

## Results

### ProChoreo Overview

The ESM2 3B model first processes the primary amino acid sequence, distilling from it a rich representation that captures both evolutionary patterns and local context. In parallel, a set of conformational snapshots, derived from MD trajectories for the same protein, provides an ensemble-level view of its structural heterogeneity. Each member of the ensemble is encoded via an Equivariant Graph Neural Network (EGNN) with nodes initialized according to the SurfPro^23^ and MaSIF ^24^ focusing on C_α_ coordinates to capture spatial relationships relevant to the protein’s folding and dynamic motions.

A contrastive learning paradigm then aligns these two data streams, associating sequence-derived features with the corresponding ensemble embeddings. This procedure produces a distilled model that translates sequence information into a latent space reflecting conformational diversity. In the subsequent design phase, the sequence embedding is fused with the ensemble embedding and fed into an autoregressive protein generator, which produces novel amino acid sequences hypothesized to maintain, or even enhance, desired structural states observed in the MD simulations. Finally, the newly generated candidates undergo *in silico* assessments of protein–protein interaction profiles, alongside further MD-based analyses of their dynamic stability (**Fig. 1**).

**Fig. 1:**
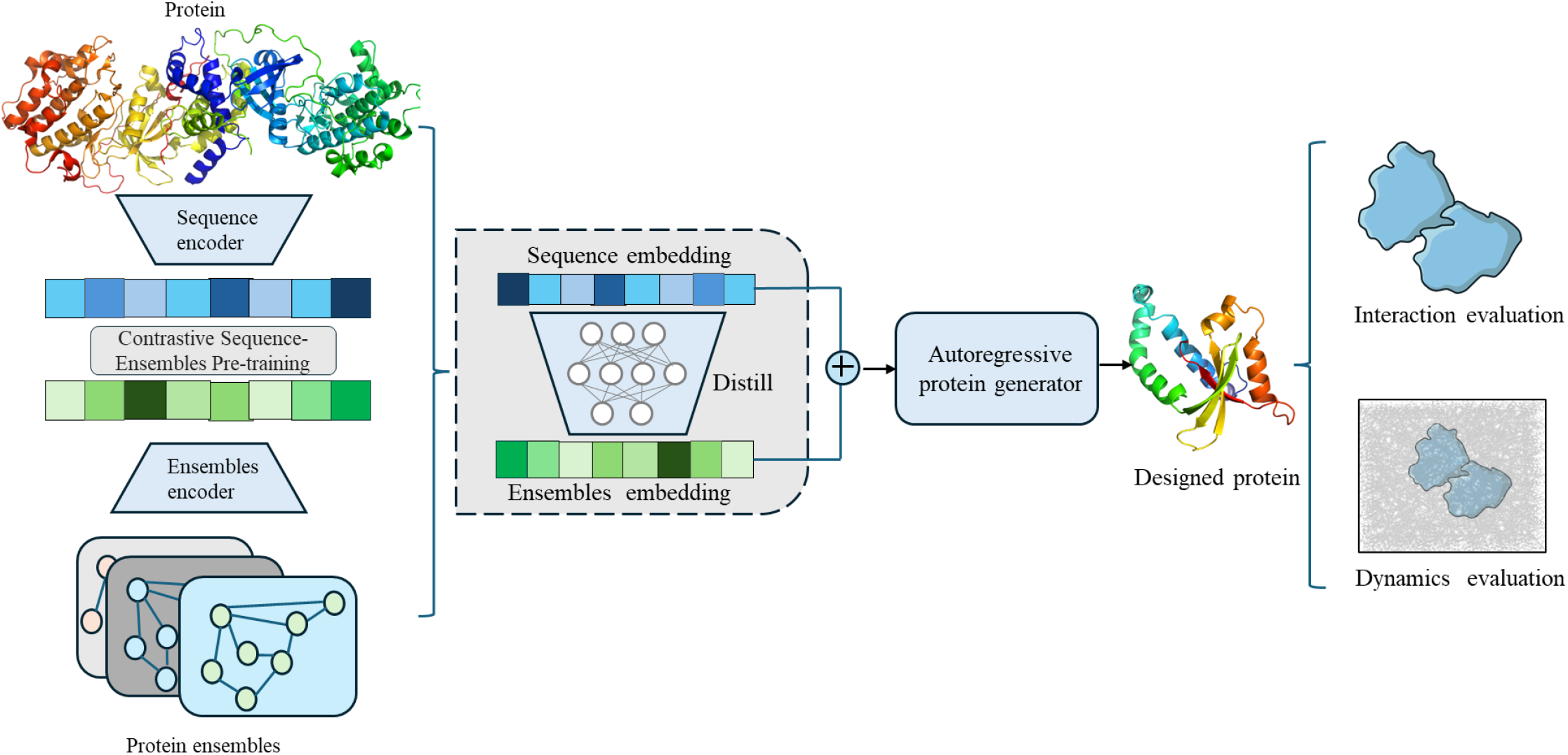
The overview of ProChoreo. The ProChoreo integrates sequence and structural ensemble representations for protein design. Sequence features are extracted using the ESM2 3B model, while MD trajectories generate protein ensembles, represented using an EGNN with Cα embeddings. A contrastive learning framework aligns sequence and ensemble embeddings, followed by a distilled model to learn sequence-to-ensemble mapping. The fused embeddings are input into an autoregressive protein generator to design new proteins. Designed proteins are validated through protein–protein interaction evaluation and MD simulations to ensure functional relevance.

### Dataset

To enable representation learning of protein conformational dynamics, we constructed a comprehensive molecular dynamics (MD) ensemble dataset consisting of both membrane proteins and non-membrane proteins. The membrane protein subset comprises 117 GPCR trajectories, each simulated for 500 ns, representing a set of receptor families with distinct signaling mechanisms and structural rearrangements (**Fig. 2 a–e**). Specifically, the dataset includes CXCR4 chemokine receptor, 5-hydroxytryptamine receptor 2B (5-HT_2_B), 5-hydroxytryptamine receptor 1B (5-HT_1_B), nociceptin/orphanin FQ receptor (NOP), and adenosine A_2_A receptor (A_2_AAR). The GPCR ensembles capture characteristic conformational transitions associated with receptor activation, such as the outward displacement of transmembrane helix 6 (TM6) in CXCR4 highlighted by the red dashed line (**Fig. 2a**), a hallmark structural indicator of the active state across Class A GPCRs. In addition, the dataset contains 4170 non-membrane protein trajectories, each simulated for 100 ns, covering diverse soluble enzymes and structural scaffolds. An example is IMPa metallopeptidase (**Fig. 2f**), which exhibits loop rearrangements and domain flexibility.

**Fig. 2:**
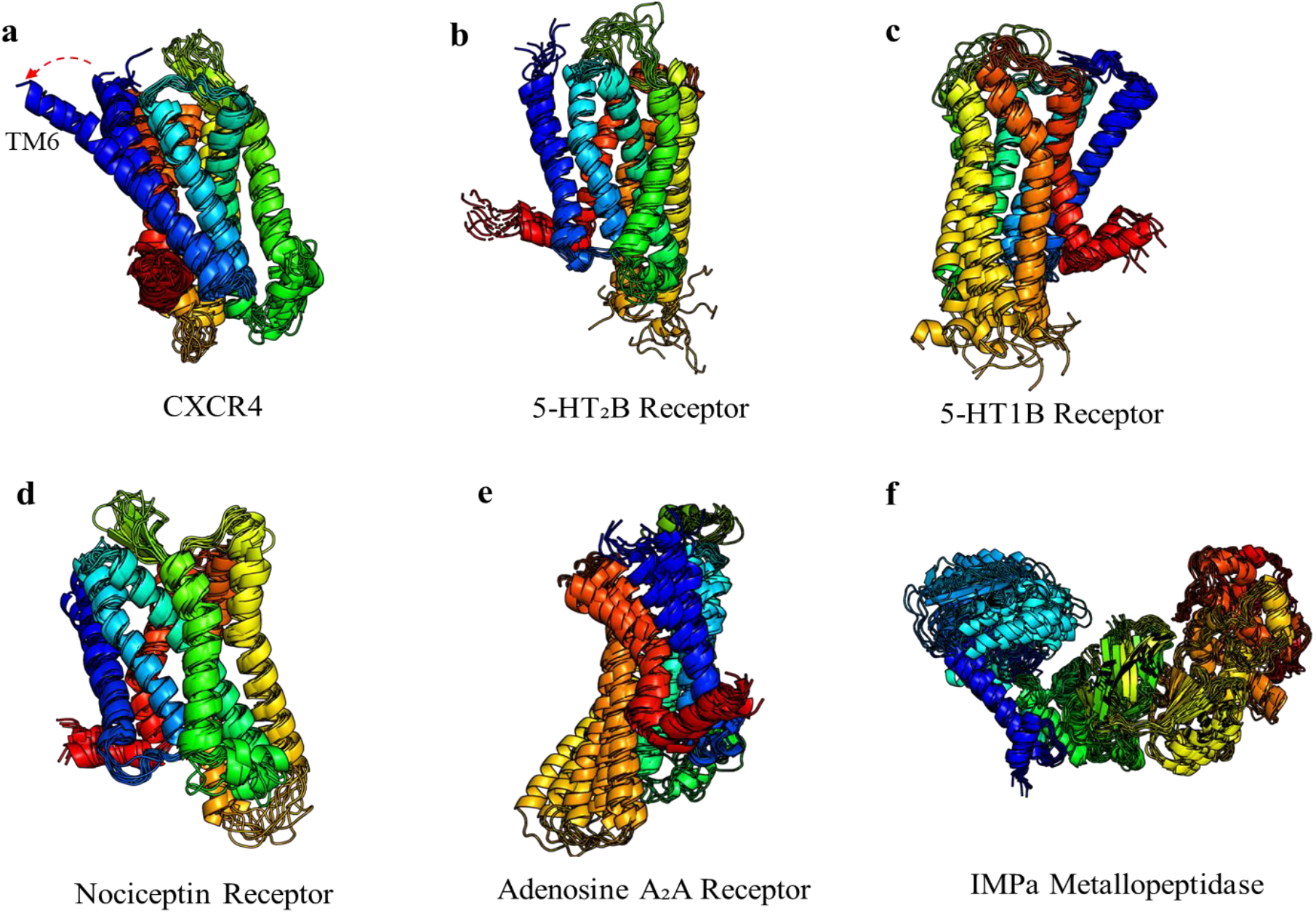
Structural ensembles of representative G protein–coupled receptors (GPCRs) (a, b, c, d, and e) and non-membrane protein (f). Protein examples shown highlight the conformational diversity among proteins. (a) CXCR4 chemokine receptor: red dashed line indicates the outward displacement of transmembrane helix 6 (TM6) observed in the active-state conformation. (b) 5-Hydroxytryptamine receptor 2B (5-HT_2_B). (c) 5-Hydroxytryptamine receptor 1B (5-HT_1_B). (d) Nociceptin/orphanin FQ receptor (NOP). (e) Adenosine A_2_A receptor (A_2_AAR). (f) IMPa metallopeptidase.

### Pretraining

We evaluated the pretrained bidirectional contrastive framework by quantifying the correspondence between protein sequence and conformational ensemble representations. The model was trained on paired data to align latent embeddings from the sequence encoder and the ensemble encoder, and evaluated in two retrieval settings, which are sequence-to-ensemble (Seq2Ens) and ensemble-to-sequence (Ens2Seq). As shown in **Fig. 3a**, both retrieval directions demonstrate consistent alignment between the two modalities. The Seq2Ens model reached R@1 = 0.7884, R@5 = 0.9299, and R@10 = 0.9580, whereas the reciprocal Ens2Seq retrieval achieved R@1 = 0.7603, R@5 = 0.9180, and R@10 = 0.9495. The pairwise squared distance distribution between the two modalities (**Fig. 3b**) showed that the majority of pairs are concentrated at distances below 0.5, revealing tight clustering and high consistency between sequence-derived and ensemble-derived embeddings. This compact and unimodal profile indicates effective latent alignment of structural and sequential information. Quantitatively, the representation exhibited an alignment score of 0.357 and uniformity of –3.298, reflecting measurable correspondence between modalities and comprehensive coverage of features across the embedding manifold.

**Fig. 3:**
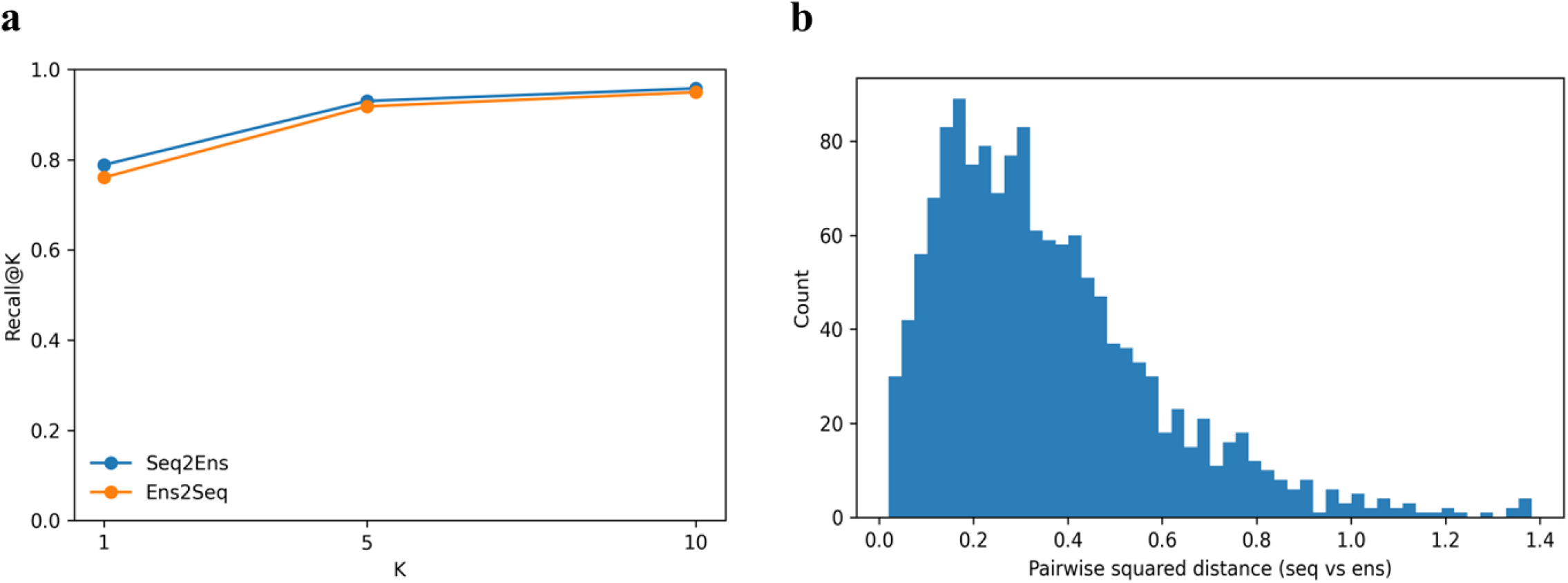
Contrastive alignment between protein sequences and conformational ensembles. (a) Bidirectional retrieval performance between protein sequence embeddings and ensemble embeddings obtained after contrastive pretraining. (b) Distribution of pairwise squared distances between the sequence and ensemble embeddings (n = 1427 pairs).

### Model Evaluation

To quantitatively assess the impact of contrastive pretraining between protein sequence and conformational ensembles, we benchmarked ProChoreo against two baselines: ProChoreo-ΔAlign, a variant trained without cross-modal alignment between sequence and ensemble representations, and PepMLM, a peptide–protein language model relying solely on sequence embeddings. Evaluation was performed across multiple receptor–binder complexes, each identified by PDB key, using a comprehensive set of structural and complex quality metrics including ptm, iptm, complex pLDDT, complex ipLDDT, complex pDE, and complex ipDE (**Table 1**).

**Table 1.** Comparison of protein–binder design performance across methods.

| PDB key | Method | confidence score | ptm | iptm | complex plddt | complex iplddt | complex pde | complex ipde |
| --- | --- | --- | --- | --- | --- | --- | --- | --- |
| 1NUN | **ProChoreo** | 0.602 ±0.004 | 0.375 ±0.008 | 0.096 ±0.020 | 0.728 ±0.006 | 0.637 ±0.036 | 1.642 ±0.094 | 13.930 ±1.353 |
| | ProChoreo-ΔAlign | 0.535 ±0.005 | 0.392 ±0.006 | 0.144 ±0.024 | 0.632 ±0.003 | 0.499 ±0.011 | 2.467 ±0.054 | 11.591 ±0.548 |
| | PepMLM | 0.586 ±0.006 | 0.389 ±0.007 | 0.126 ±0.019 | 0.701 ±0.009 | 0.591 ±0.030 | 1.592 ±0.119 | 11.411 ±0.618 |
| 3UG2 | **ProChoreo** | 0.707 ±0.014 | 0.745 ±0.011 | 0.562 ±0.044 | 0.744 ±0.007 | 0.644 ±0.013 | 1.638 ±0.078 | 10.006 ±0.814 |
| | ProChoreo-ΔAlign | 0.623 ±0.004 | 0.713 ±0.014 | 0.172 ±0.053 | 0.735 ±0.017 | 0.570 ±0.062 | 1.763 ±0.415 | 12.622 ±5.184 |
| | PepMLM | 0.670 ±0.005 | 0.688 ±0.002 | 0.108 ±0.012 | 0.810 ±0.004 | 0.792 ±0.019 | 1.275 ±0.033 | 16.504 ±0.906 |
| 4HSA | **ProChoreo** | 0.621 ±0.011 | 0.605 ±0.011 | 0.188 ±0.029 | 0.730 ±0.012 | 0.586 ±0.029 | 1.398 ±0.235 | 11.365 ±1.784 |
| | ProChoreo-ΔAlign | 0.583 ±0.003 | 0.589 ±0.008 | 0.114 ±0.016 | 0.701 ±0.005 | 0.543 ±0.031 | 1.735 ±0.182 | 14.381 ±1.492 |
| | PepMLM | 0.606 ±0.001 | 0.598 ±0.005 | 0.150 ±0.034 | 0.720 ±0.008 | 0.559 ±0.044 | 1.480 ±0.146 | 12.795 ±2.352 |
| 4I23 | **ProChoreo** | 0.755 ±0.003 | 0.714 ±0.011 | 0.706 ±0.020 | 0.768 ±0.006 | 0.672 ±0.027 | 1.743 ±0.090 | 11.888 ±1.114 |
| | ProChoreo-ΔAlign | 0.587 ±0.006 | 0.676 ±0.005 | 0.120 ±0.013 | 0.704 ±0.008 | 0.626 ±0.019 | 2.708 ±0.067 | 18.285 ±0.796 |
| | PepMLM | 0.734 ±0.013 | 0.725 ±0.008 | 0.720 ±0.044 | 0.737 ±0.006 | 0.639 ±0.018 | 1.484 ±0.096 | 9.263 ±1.155 |
| 5IUS | **ProChoreo** | 0.635 ±0.016 | 0.576 ±0.056 | 0.239 ±0.148 | 0.734 ±0.017 | 0.668 ±0.018 | 1.608 ±0.142 | 16.608 ±6.292 |
| | ProChoreo-ΔAlign | 0.556 ±0.004 | 0.546 ±0.018 | 0.106 ±0.039 | 0.668 ±0.010 | 0.556 ±0.037 | 1.965 ±0.103 | 14.918 ±3.134 |
| | PepMLM | 0.599 ±0.009 | 0.569 ±0.021 | 0.163 ±0.029 | 0.708 ±0.015 | 0.607 ±0.013 | 1.460 ±0.074 | 11.123 ±2.459 |
| 5MZV | **ProChoreo** | 0.689 ±0.014 | 0.632 ±0.014 | 0.193 ±0.056 | 0.813 ±0.005 | 0.701 ±0.029 | 1.372 ±0.053 | 13.478 ±1.615 |
| | ProChoreo-ΔAlign | 0.612 ±0.005 | 0.628 ±0.005 | 0.130 ±0.003 | 0.733 ±0.006 | 0.572 ±0.033 | 1.666 ±0.071 | 12.043 ±0.414 |
| | PepMLM | 0.665 ±0.002 | 0.642 ±0.012 | 0.180 ±0.022 | 0.786 ±0.004 | 0.658 ±0.031 | 1.179 ±0.028 | 8.934 ±0.706 |
| 6LUD | **ProChoreo** | 0.671 ±0.014 | 0.730 ±0.015 | 0.225 ±0.054 | 0.783 ±0.006 | 0.635 ±0.023 | 1.555 ±0.222 | 10.654 ±2.140 |
| | ProChoreo-ΔAlign | 0.623 ±0.004 | 0.712 ±0.011 | 0.116 ±0.040 | 0.749 ±0.006 | 0.630 ±0.027 | 1.939 ±0.043 | 14.271 ±2.086 |
| | PepMLM | 0.651 ±0.005 | 0.707 ±0.013 | 0.151 ±0.039 | 0.776 ±0.006 | 0.611 ±0.047 | 1.603 ±0.222 | 13.588 ±3.105 |
| 7KNU | **ProChoreo** | 0.693 ±0.010 | 0.671 ±0.010 | 0.484 ±0.040 | 0.745 ±0.003 | 0.633 ±0.006 | 1.736 ±0.104 | 12.032 ±0.760 |
| | ProChoreo-ΔAlign | 0.574 ±0.005 | 0.610 ±0.007 | 0.122 ±0.016 | 0.687 ±0.004 | 0.544 ±0.032 | 2.576 ±0.217 | 19.454 ±0.896 |
| | PepMLM | 0.623 ±0.032 | 0.646 ±0.030 | 0.251 ±0.128 | 0.715 ±0.008 | 0.597 ±0.022 | 2.348 ±0.274 | 16.582 ±1.751 |
| 7KPB | **ProChoreo** | 0.566 ±0.008 | 0.526 ±0.012 | 0.171 ±0.027 | 0.664 ±0.011 | 0.590 ±0.039 | 1.523 ±0.098 | 8.928 ±0.366 |
| | ProChoreo-ΔAlign | 0.495 ±0.002 | 0.518 ±0.010 | 0.147 ±0.012 | 0.581 ±0.004 | 0.517 ±0.021 | 2.565 ±0.115 | 10.215 ±1.157 |
| | PepMLM | 0.553 ±0.004 | 0.522 ±0.012 | 0.162 ±0.012 | 0.651 ±0.006 | 0.581 ±0.022 | 1.596 ±0.038 | 9.395 ±0.344 |
| 7PCD | **ProChoreo** | 0.635 ±0.005 | 0.721 ±0.008 | 0.210 ±0.024 | 0.741 ±0.004 | 0.606 ±0.019 | 1.294 ±0.027 | 8.895 ±0.603 |
| | ProChoreo-ΔAlign | 0.566 ±0.003 | 0.694 ±0.004 | 0.143 ±0.025 | 0.672 ±0.008 | 0.542 ±0.015 | 2.286 ±0.031 | 13.010 ±0.451 |
| | PepMLM | 0.610 ±0.012 | 0.713 ±0.006 | 0.233 ±0.057 | 0.704 ±0.004 | 0.590 ±0.019 | 2.129 ±0.072 | 12.046 ±0.753 |
| 7RG9 | **ProChoreo** | 0.754 ±0.017 | 0.732 ±0.019 | 0.729 ±0.037 | 0.760 ±0.012 | 0.722 ±0.021 | 1.664 ±0.236 | 10.139 ±3.593 |
| | ProChoreo-ΔAlign | 0.604 ±0.006 | 0.692 ±0.010 | 0.160 ±0.029 | 0.715 ±0.011 | 0.587 ±0.017 | 2.094 ±0.433 | 14.679 ±3.739 |
| | PepMLM | 0.705 ±0.016 | 0.754 ±0.017 | 0.597 ±0.037 | 0.732 ±0.016 | 0.687 ±0.042 | 1.418 ±0.322 | 5.504 ±2.363 |
| 7V3R | **ProChoreo** | 0.611 ±0.010 | 0.678 ±0.008 | 0.094 ±0.017 | 0.740 ±0.008 | 0.619 ±0.033 | 2.069 ±0.124 | 16.455 ±1.839 |
| | ProChoreo-ΔAlign | 0.571 ±0.003 | 0.671 ±0.006 | 0.095 ±0.008 | 0.690 ±0.003 | 0.596 ±0.041 | 2.326 ±0.083 | 14.486 ±0.722 |
| | PepMLM | 0.592 ±0.005 | 0.672 ±0.011 | 0.120 ±0.026 | 0.710 ±0.008 | 0.562 ±0.029 | 1.894 ±0.178 | 13.157 ±2.187 |

Across all test systems, ProChoreo consistently outperformed both baselines in confidence-score and structural fidelity metrics. The model achieved an average confidence-score improvement of 4–8% over ProChoreo-ΔAlign and up to 12% over PepMLM. The gains were particularly pronounced in global structure agreement (ptm and iptm) and complex-level pLDDT, underscoring the contribution of ensemble-conditioned representations to capturing realistic interfacial geometry. For example, in complexes such as 4I23, 6LUD, and 7KNU, ProChoreo achieved complex pLDDT values exceeding 0.74 with correspondingly lower predicted energy, indicating improved structural confidence and thermodynamic plausibility.

The inclusion of the alignment module between sequence and ensemble embeddings was critical for this performance. Removing it (ProChoreo-ΔAlign) resulted in systematic drops in iptm and complex ipDE that are metrics that reflect interface precision, demonstrating that alignment learning enables the model to better reconcile dynamic conformational information with sequence-level representations. In contrast, the PepMLM baseline, though pretrained on large-scale sequence corpora, lacked explicit ensemble conditioning and consequently exhibited reduced complex pLDDT and elevated pDE across all targets, indicating poorer folding-interface compatibility.

### Case study

To evaluate the capacity of ProChoreo to generate functionally competent binders, we first examined the interaction between the designed binder produced by our model and selected based on structural prediction and interaction assessment using Boltz 1 and the human sweet taste receptor subunit TAS1R2, a C family of GPCR primarily responsible for recognizing sugars and high-potency sweeteners. For comparison, we simulated the complex between TAS1R2 and the natural sweet protein brazzein, which serves as a validated reference ligand (**Fig. 4a**). Both complexes were subjected to 500 ns MD simulations, and the resulting trajectories were analyzed using MM-GBSA to estimate binding free energies. The calculated affinities were –50.79 kJ mol^−1^ for the TAS1R2–binder complex and –69.99 kJ mol^−1^ for TAS1R2–brazzein, indicating that brazzein exhibits stronger binding than designed binder (**Fig. 4b**, left). Interestingly, this observation is consistent with prior reports that natural sugars display markedly weaker affinity toward TAS1R2.

**Fig. 4:**
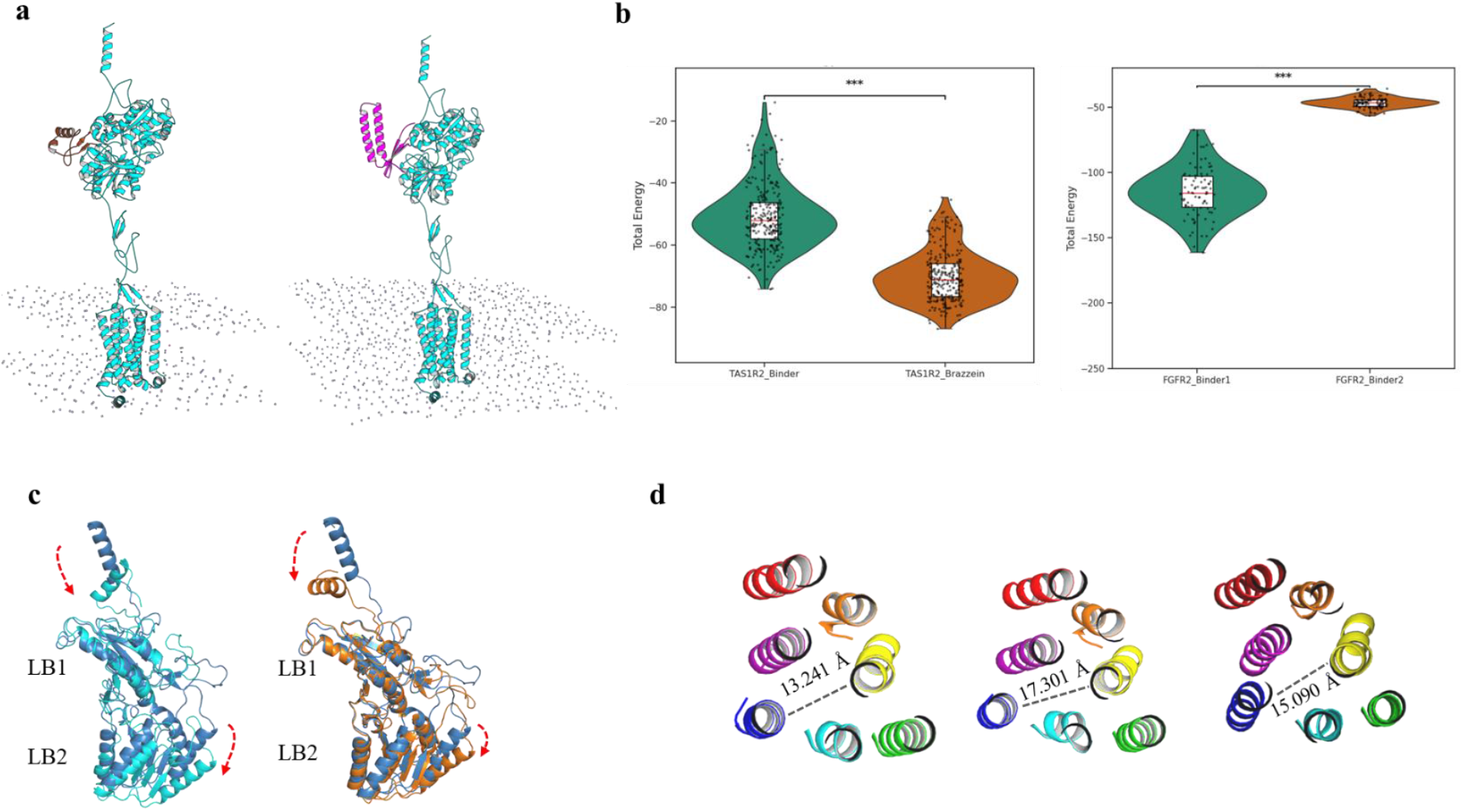
Case studies demonstrating binder design and conformational analysis of representative receptors. (a) Molecular dynamics (MD) simulation snapshots of receptor–binder complexes for the human sweet taste receptor (TAS1R2, left) and FGFR2 (right), showing stable binding configurations within the membrane environment. (b) Total energy distributions from MD trajectories comparing ProChoreo-designed binders with natural counterparts (brazzein for TAS1R2 and native ligand for FGFR2); *** indicates p < 0.001. (c) Venus flytrap (VFT) domain of TAS1R2 showing the relative motion of LB1 and LB2 lobes upon binder engagement. (d) Top-down view of TAS1R2 VFT domain illustrating inter-lobe distances across apo (left), brazzein-bound (middle), and binder-bound states (right).

We next analyzed the conformational dynamics of TAS1R2 during complex formation. In both the designed binder and brazzein complexes, the Venus Flytrap Domain (VFTD) exhibited a characteristic closure of lobe 1 (LB1) accompanied by a modest opening of lobe 2 (LB2) (**Fig. 4c**), consistent with activation-associated rearrangements reported for other class C GPCRs. Examination of the transmembrane (TM) helices further revealed similar allosteric signatures reflected by coordinated outward displacement of TM3 and TM6^25^. The distance between TM3 and TM6 increased from 13.24 Å in the apo state to 15.09 Å in the binder complex and 17.30 Å in the brazzein complex (**Fig. 4d**), reflecting an outward displacement of TM6 upon ligand engagement. These results collectively suggest that the designed binder, while exhibiting weaker binding than brazzein, stabilizes an active-like conformation of TAS1R2 reminiscent of natural sweet protein interactions.

In another case study, we applied ProChoreo to design binders targeting the receptor tyrosine kinase FGFR2, representing a structurally and functionally distinct receptor class. The designed FGFR2 binder underwent 100 ns MD simulation, during which the complex remained stable throughout the trajectory (**Fig. 4b**, right). The MM-GBSA–derived binding free energy exceeded –100 kJ mol^−1^, demonstrating strong and persistent interaction with the receptor interface. These results confirm that ProChoreo can design binders with robust interaction propensities across both GPCR and RTK families.

## Discussion

Our results demonstrate that ProChoreo successfully extends sequence-based protein generation into the dynamic regime by explicitly incorporating conformational ensemble information. In the case of the human sweet taste receptor subunit TAS1R2, the designed binder reproduced several hallmarks of receptor activation observed in the natural sweet protein brazzein, including the closure of LB1 and the partial opening of LB2 within the VFTD. The outward displacement of TM6 from 13.24 Å in the apo form to 15.09 Å in the binder complex, and to 17.30 Å with brazzein, mirrors the allosteric movement associated with GPCR activation, suggesting that the designed binder stabilizes an active-like conformation of TAS1R2 despite its moderately weaker binding affinity (–50.79 kJ mol^−1^ vs. –69.99 kJ mol^−1^ for brazzein). This observation highlights that binding energetics alone do not fully determine functional engagement, and that ensemble-informed design can capture subtle conformational cues underlying receptor modulation.

The analysis of the FGFR2 system further supports the generalizability of our approach across receptor classes. The designed FGFR2 binder maintained structural stability over 100 ns MD simulation and exhibited strong binding as –100 kJ mol^−1^, consistent with tight interface complementarity predicted during generation. Together, these findings indicate that incorporating ensemble-aware representations allows the model to infer energetically favorable and dynamically coherent interaction geometries across mechanistically distinct receptor families.

Nevertheless, the current framework remains constrained by the quality and diversity of molecular dynamics–derived ensembles, which may not fully capture the conformational space sampled under experimental conditions. In future work, we aim to integrate experimental ensemble data, such as NMR and cryo-EM variability analyses, and learned diffusion-based conformational models to expand coverage of the protein energy landscape. Extending ProChoreo toward direct sequence-to-ensemble generation represents a natural next step toward designing proteins that not only bind tightly but also modulate structural dynamics in a functionally programmable manner.

## Methods

### MD dataset

The ATLAS database (https://www.dsimb.inserm.fr/ATLAS) was employed to obtain all-atom MD simulations of a diverse range of soluble protein structures. The database comprises 1390 soluble protein chains, each simulated for 100 ns across 3 independent replicates^26^. To further examine receptor dynamics, an additional GPCRmd dataset (http://gpcrmd.org/) consisting of 39 systems, each simulated for 500 ns across 3 replicates was incorporated^27^. In total, the combined MD dataset includes 1429 systems, representing a cumulative simulation time of 475.5 µs.

### Protein conformational ensembles generation

MD trajectories were analyzed using the MDAnalysis toolkit to extract representative conformational states from each protein system^28^. For each system, trajectory and topology files were merged using MDAnalysis, and the backbone atoms were aligned to minimize structural variability. Principal component analysis (PCA) was then performed on the aligned backbone atoms, retaining the first five components (n_components = 5) to capture the dominant modes of motion. The resulting PCA projections were subjected to KMeans clustering (n_clusters = 10), partitioning the combined replicate data into ten distinct conformational states. For each cluster, the frame with the minimal distance to the cluster centroid was selected as the representative state, and the Cα atom coordinates were extracted^28,29^.

### Protein design dataset

For protein design applications, we incorporated a dataset derived from DIPS^30,31^. Only pairs meeting the following conditions were retained: (i) the molecule is a protein containing at least 50 amino acids; (ii) the structure was determined by Cryo-EM or X-ray crystallography at a resolution superior to 3.5 Å; (iii) the buried surface area upon binding exceeds 500 Å^2^. After filtering, a total of 134,215 chain pairs were retained.

### Seq2Ens-CLIP Module

In this module, we developed a contrastive learning framework, inspired by the CLIP architecture, to align protein sequence embeddings with geometric ensemble representations^21^. The protein sequence encoder was instantiated using the pre-trained ESM2_t36_3B_UR50D model^8^, which provided residue-level embeddings. In parallel, a geometric encoder based on an EGNN was employed to capture the structural information of proteins, using Cα atoms as the nodes of the graph^32^. The raw embeddings generated by the sequence encoder and the EGNN were denoted as *seq*-*emb* ∈ *R*^*B*×*N*×*d*^ and *geom*-*emb* ∈ *R*^*B*×*N*×*d*^, respectively, where B is the batch size, N is the number of residues per protein, and d is the embedding dimension.

To measure the similarity between these embeddings, they were L2-normalized and projected to same dimensions. The similarity matrix (*logits* ∈ *R*^*M*×*M*^) was then computed as the dot product of the normalized embeddings, where *M* represents the total number of valid residue positions.

The sequence-to-geometry loss was defined as l_seq_ = CrossEntropy(*logits, y*) and the geometry-to-sequence loss as l_geom_ = CrossEntropy(*logits*^T^, *y*), where (*y* = [0,1, …, *M* − 1]). The final loss was computed in both the sequence-to-geometry and geometry-to-sequence directions and defined as: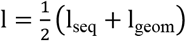. A following model^16^ further distilled a mapping from seq-emb to geom-emb.

### Protein Generator Module

We propose a generative method that produces a complementary protein chain *B* conditioned on a target chain *A*. Chain A is provided as an amino acid sequence *x*_*A*_ = (*x*_1_, *x*_2_, …, *x*_*L*_). In our approach, chain *A* is first tokenized and padded (or truncated) to a fixed length, and then processed by a large-scale pretrained language model (ESM2_t36_3B_UR50D) to extract high-dimensional, residue-level embeddings. A binary mask is generated concurrently to indicate the valid positions in the sequence. These sequence embeddings are further refined by a geometry-distilled network (ProClipStudent), which computes structure-aware features from the original embeddings. Because the geometry-derived features have a lower dimensionality, a trainable linear projection is applied to map them into the same space as the ESM embeddings. The projected geometry features are then combined with the original embeddings via elementwise addition to form a fused representation that captures both contextual and structural information. This fused representation is used as the memory input for a Transformer-based autoregressive decoder that generates chain *B* one residue at a time. *y*_*t*_ is the token predicted at time step t (with *V* denoting the vocabulary of valid amino acids) During training, the decoder minimizes a cross-entropy loss between the predicted tokens and the ground-truth sequence, while during inference the model generates tokens either by greedy selection or by sampling from the softmax distribution (with optional temperature scaling and top-*k* filtering). The entire pipeline can be summarized by the following formula:

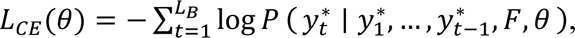

Where *x*_*A*_ denotes the input protein chain *A*.

*E*_*ESM*_(*x*_*A*_) represents the ESM-based embeddings extracted from chain *A*.

Student(·) computes the geometry-distilled embedding from *E*_*ESM*_(*x*_*A*_).

*W* is a trainable projection matrix that aligns the geometry-derived features with the ESM embeddings.

*y*_*t*_ is the token predicted at time step *t* (with *V* denoting the vocabulary of valid amino acids).

*F* is the fused representation used as memory in the decoder.

*θ* represents the parameters that are updated during training.

### Training

During training, our model was optimized using the Adam optimizer with an initial learning rate of 1 × 10^−4^ and a weight decay of 1 × 10^−4^. The learning rate was scheduled via a cosine annealing strategy, implemented with PyTorch’s CosineAnnealingLR. To stabilize training and reduce memory consumption, mixed-precision training was enabled using PyTorch’s GradScaler. The training was conducted on two Nvidia A100 40 GB GPUs.

### Eval

Protein–protein interactions were evaluated using Boltz-1^22^, a substitute method for AlphaFold3^33^, to assess the complementarity of the binding interfaces between the generated proteins and the target protein sequence. To further investigate the structural stability and dynamic behavior of the predicted complexes, MD simulations were performed using the GROMACS^34^ software package (version 2022) with the CHARMM36 force field^35^.

### Claim

Computations for this research were performed on the Pennsylvania State University’s Institute for Computational and Data Sciences’ Roar Collab supercomputer.

### Data availability

The data generated in this study will be made publicly available upon publication. Experimental data related to this work are available from the corresponding author upon reasonable request.

### Code

https://github.com/SaaaaiDing/PD_NLP

## Acknowledgements

This work was supported by Rasmussen Endowment by The Pennsylvania State University, as well as the USDA National Institute of Food and Agriculture and Hatch Appropriations under Accession number 7005076.

## Author contributions

Conceptualization: Y.Z. and S.D. Methodology: S.D. and Y.Z. Software: S.D. Formal analysis: S.D. Investigation: S.D. Visualization: S.D. Validation: S.D. and Y.Z. Writing-original draft: S.D. and Y.Z. Writing-review and editing: S.D. and Y.Z. Supervision: Y.Z.

